# *In vitro* and intracellular activity of imipenem combined with tedizolid, rifabutin, and avibactam against *Mycobacterium abscessus*

**DOI:** 10.1101/411181

**Authors:** Eva Le Run, Michel Arthur, Jean-Luc Mainardi

## Abstract

*Mycobacterium abscessus* has emerged as a significant pathogen responsible for chronic pulmonary infections in cystic fibrosis (CF) patients, which are difficult to treat due to resistance to a broad range of antibiotics. The initial phase of the recommended treatment in CF patients includes imipenem used without any β-lactamase inhibitor in spite of the production of the β-lactamase Bla_Mab_. Here, we determine whether the addition of tedizolid, a once-daily oxazolidinone, improves the activity of imipenem alone or in combination with a β-lactamase inhibitor, avibactam, and rifabutin.

The activity of the drugs was evaluated against *M*. *abscessus* CIP104536 by determining *in vitro* and intracellular antibacterial activities. The impact of Bla_Mab_ inhibition by avibactam on antibiotic activity was assessed by comparing CIP104536 and its β-lactamase-deficient derivative (Δ*bla*_Mab_).

The minimal inhibitory concentrations (MICs) of tedizolid against *M*. *abscessus* CIP104536 and Δ*bla*_Mab_ were 4 μg/mL. Tedizolid combined with imipenem showed a moderate synergistic effect with fractional inhibitory concentration (FIC) indexes of 0.41 and 0.38 for CIP104536 and Δ*bla*_Mab_, respectively. For both strains, the addition of tedizolid at 2 μg/mL, corresponding to the peak serum concentration, increased the intracellular efficacy of imipenem at 8 and 32 μg/mL. Addition of avibactam and rifabutin improved the activity of the imipenem-tedizolid combination against CIP104536S.

The imipenem-tedizolid combination should be further considered for the treatment of *M. abscessus* pulmonary infections in CF patients. The efficacy of the treatment might benefit from the use of a β-lactamase inhibitor, such as avibactam, and the addition of rifabutin.

---

Nontuberculous mycobacteria (NTM) have been increasingly isolated from the lungs of cystic fibrosis (CF) patients (1–4) with a prevalence estimated at 11 to 13% (1,5). Among NTM, *M*. *abscessus,* a rapidly growing mycobacterium, is the predominant species isolated in these patients (40 to 50%) (2,6). Chronic pulmonary *M*. *abscessus* infections have been associated with increased lung function decline in CF patients (5).

The treatment is particularly complex and difficult since *M. abscessus* is intrinsically resistant to a broad range of antibiotics, including those used for the treatment of tuberculosis (7,8). In 2016, the US Cystic Fibrosis Foundation and European Cystic Fibrosis Society have established recommendations for the management of NTM in CF patients (9). The typical treatment schedule consists of an initial phase with the combination of a carbapenem (imipenem), a macrolide (azithromycin), an aminoglycoside (amikacin), and a glycylcycline (tigecycline) for a duration of 3 months (9). Four *per os* drugs (azithromycin, minocycline, clofazimine, and moxifloxacin) are proposed for the continuation phase for at least 12 months (9). In spite of these lengthy courses of antibiotics, the prognosis of pulmonary infections is poor in the context of CF with a cure rate of 30 to 50% (10,11). In case of macrolide resistance, present in 40 to 60% of the isolates (12), the rate of bacteriological eradication is in the order of 25% (10). In this context, there is an urgent need to identify additional therapeutic options. This could be achieved in the short term by repurposing existing drugs approved for the treatment of other bacterial infections.

*M. abscessus* isolates produce a broad-spectrum β-lactamase Bla_Mab_, which hydrolyzes most β-lactams, except cefoxitin, and inactivates imipenem at a very slow rate (13,14). Imipenem is currently used in the absence of any β-lactamase inhibitor although Bla_Mab_ was recently shown to limit the intracellular activity of imipenem in human macrophages (13,15,16). First generation of β-lactamase inhibitors, clavulanate, tazobactam, and sulbactam, are inactive against Bla_Mab_ (17). However, Bla_Mab_ is inhibited by a novel β- lactamase inhibitor, avibactam (13), which has been developed in combination with ceftazidime for the treatment of infections due to multi-drug resistant Enterobacteriaceae (18). Avibactam extends the spectrum of β-lactams active against *M*. *abscessus* (13,15) and improves the efficacy of imipenem, both in macrophages and in zebrafish embryos (16).

Linezolid, a first-generation oxazolidinone antibiotic, is an important therapeutic option for infections caused by resistant Gram-positive bacteria (19). In *Mycobacterium tuberculosis*, clinical efficacy of linezolid has been reported in most difficult-to-treat MDR and XDR-cases (20,21). Linezolid has been positioned by WHO in the group 5 of anti-TB drugs for the treatment of MDR and XDR-TB cases (22). In case of intolerance to drugs of the reference treatment for *M*. *abscessus* in CF patients, linezolid is also proposed during the continuation phase (9), in spite of high minimal inhibitory concentrations (MICs) comprised between 16 and 64 μg/mL (7,23,24). However, the high frequency and severity of adverse events limit the long-term use of linezolid (20). Tedizolid is a recently developed once-daily oxazolidinone antibiotic, which has been approved for the treatment of acute bacterial skin and soft tissue infections (25). In *M*. *abscessus,* a recent *in vitro* study has shown that tedizolid has a better *in vitro* activity compared to linezolid, as the MIC_50_s and MIC_90_s of tedizolid (2 and 8 μg/mL, respectively) were 2- to 16-fold lower than those of linezolid (24).

In this study, we have investigated the interest of repurposing tedizolid for *M. abscessus* infections in combination with imipenem alone or with imipenem, avibactam and rifabutin. We report the *in vitro* and intracellular antibacterial activities of various combinations of these four drugs. The impact of β-lactamase production was assessed by comparing *M. abscessus* CIP104536 and a derivative obtained by deletion of the gene encoding Bla_Mab_.

## RESULTS

### MICs of tedizolid and imipenem and synergy between these drugs

The MIC of tedizolid was 4 μg/mL against both *M*. *abscessus* CIP104536 and its β-lactamase deficient (Δ*bla*_Mab_) derivative (Table 1). The MICs of imipenem were 4 μg/mL and 2 μg/mL against CIP104536 and its Δ*bla*_Mab_ derivative. Against CIP104536 (Table 1), the combination of tedizolid with imipenem showed a moderate synergistic effect with a FIC index of 0.41 at 1/4 of the MIC of imipenem and 1/8 of the MIC of tedizolid. Against the Δ*bla*_Mab_ derivative, the FIC index was 0.38 at 1/4 of the MIC of imipenem and 1/8 of the MIC of tedizolid.

**Table 1.** Activity of imipenem and tedizolid against *M. abscessus* FIC-fractional inhibitory concentration index; IPM-imipenem; MIC-minimal inhibitory concentration; TZD-tedizolid

| Strain of *M. abscessus* | MIC (µg/mL) |  | FIC index |
| --- | --- | --- | --- |
|  | TZD | IPM | TZD + IPM |
| CIP104536 | 4 | 4 | 0.41 |
| CIP104536Δ*bla*<sub>Mab</sub> | 4 | 2 | 0.38 |

### In vitro killing of M. abscessus by tedizolid alone, in combination with imipenem, or in combination with imipenem and avibactam

Tedizolid was tested at ½ fold the MIC (2μg/mL) corresponding to the peak serum concentration for an administration of 200 mg per day (27) and imipenem at 2 fold the MIC (8 μg/mL). Against CIP104536, tedizolid alone had no effect in the reduction of CFUs (Fig. 1A and Table S1). A 0.7 Log_10_-reduction of CFUs was observed for imipenem alone at 8 μg/mL. The addition of tedizolid did not increase the bacterial killing for imipenem (1.0 Log_10_-reduction of CFUs). Similar results were obtained for Δ*bla*_Mab_ derivative of CIP104536 (Fig. 1B and Table S2). Since the addition of tedizolid to imipenem did not improve the activity of imipenem against *M*. *abscessus* CIP104536, we investigated the benefit of adding avibactam (4 μg/mL) (Fig. 1C and Table S3). Avibactam did not potentiate the killing by imipenem alone. The triple combination of imipenem-tedizolid-avibactam was not more active than the imipenem-tedizolid combination.

**FIG 1.**
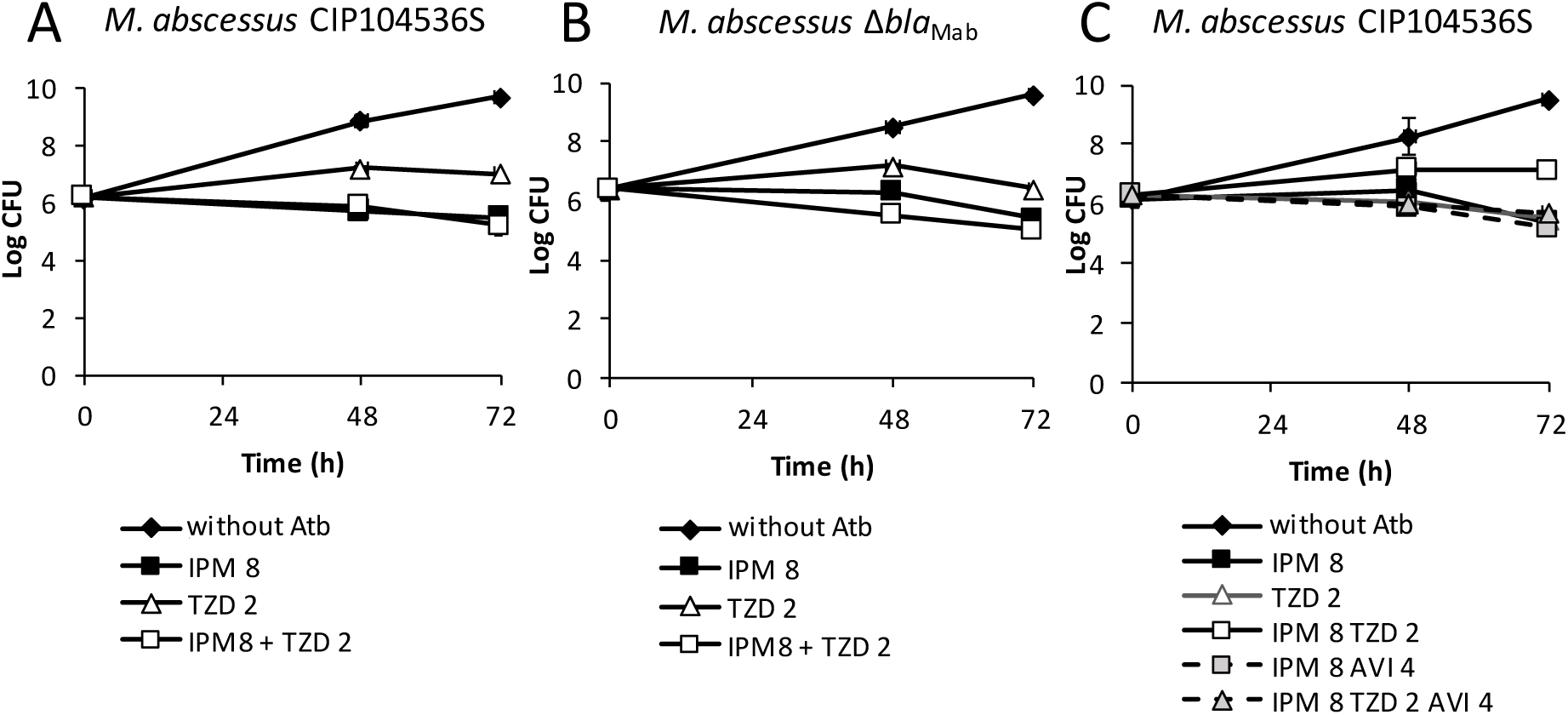
Bactericidal activity of tedizolid (TZD) alone or in combination with imipenem (IPM) and the β-lactamase inhibitor avibactam (AVI) against *M. abscessus* CIP1045636 and its Δ*bla*_Mab_ derivative. A) and B) Time kill curves of TZD at 2 μg/mL alone or in combination with IPM (8 μg/mL) against CIP104536 and its Δ*bla*_Mab_ derivative, respectively. C) Time-kill curves of TZD and IPM with or without AVI at 4 μg/mL against CIP104536. Without Atb-without antibiotic.

To conclude on the *in vitro* activity of the drugs, tedizolid did not increase the activity of imipenem against *M. abscessus*. The activity of imipenem or imipenem-tedizolid is not improved by the addition of avibactam. None of the combination was bactericidal against both strains.

### Intramacrophage activity of tedizolid alone, in combination with imipenem, or in combination with imipenem and avibactam

In the absence of antibiotic, *M*. *abscessus* CIP104536 grew in the macrophages, leading to a 143-fold increase in the number of CFUs in 48 h (Fig. 2A, Table S4). Tedizolid at 2 μg/mL partially prevented intramacrophage growth (6.85-*versus* 143-fold increase in the number of CFUs; *P* < 0.05). Imipenem at 8 μg/mL was more active than tedizolid (1.75-*versus* 6.85-fold increase in CFUs; *P* < 0.05). Increasing the concentration of imipenem from 8 μg/mL to 32 μg/mL increased the activity of this drug but the difference was not statically significant (1.75-*versus* 0.74-fold change in CFUs; *P =* 0.13). The combination of tedizolid and imipenem (8 μg/mL) was more active than imipenem alone (0.10-*versus* 1.75-fold change in CFUs; *P* < 0.05). Of note, the combination of imipenem (8 μg/mL) and tedizolid (2 μg/mL) achieved 90% killing of *M. abscessus* in the macrophage. Increasing the concentration of imipenem from 8 μg/mL to 32 μg/mL did not improve killing (90% *versus* 79%; *P* = 0.13).

**FIG 2.**
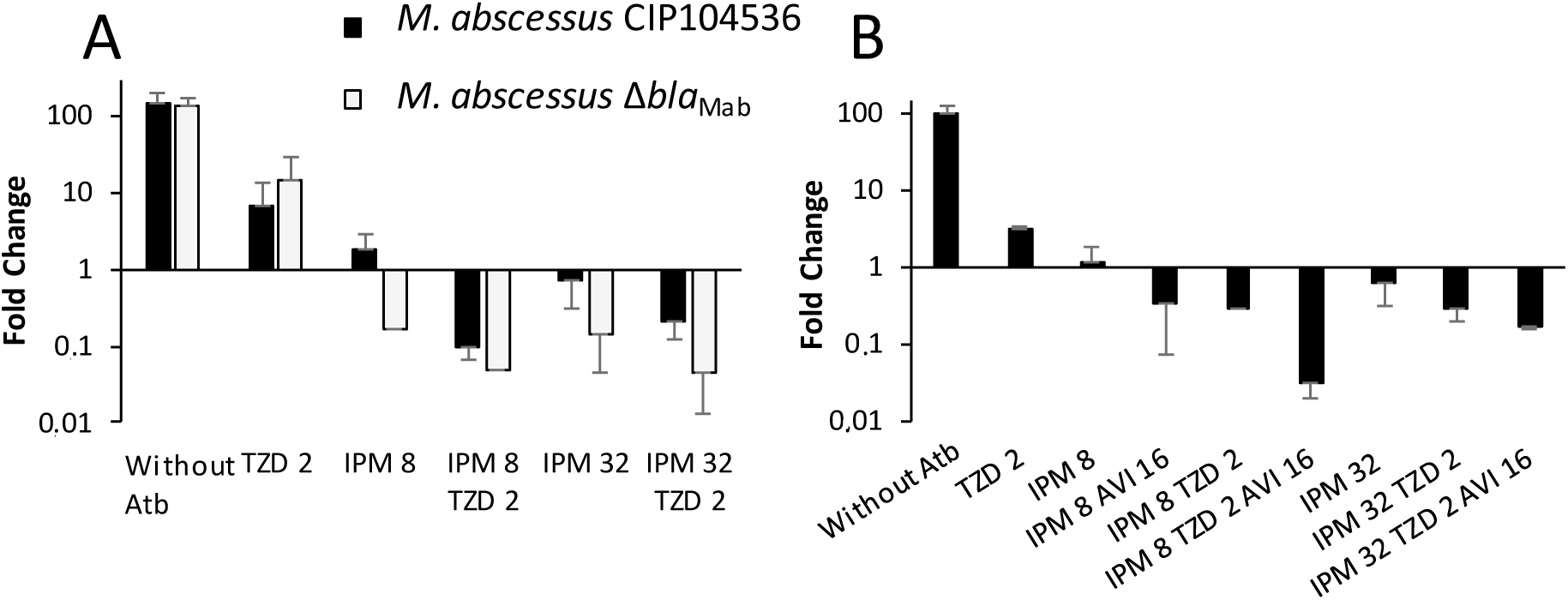
Intracellular activities of antibiotics against *M. abscessus* CIP104536 and its Δ*bla*_Mab_ derivative in the macrophage model. A) Tedizolid (TZD) at 2 μg/mL and imipenem (IPM) at 8 or 32 μg/mL were tested alone or in combination against *M. abscessus* CIP104536 and its Δ*bla*_Mab_ derivative. B) IPM at 8 and 32 μg/mL and TZD at 2 μg/mL were tested with or without AVI at 16 μg/mL against *M. abscessus* CIP104536. Without Atb-without antibiotic.

The combinations of imipenem and tedizolid were also tested against the isogenic strain *M. abscessus* CIP104536 Δ*bla*_Mab_ to evaluate the impact of the production of the β- lactamase Bla_Mab_ on the intracellular activity of the drugs (Fig. 2A and Table S5). Tedizolid alone displayed similar activity against Δ*bla*_Mab_ and CIP104536 (14.27-*versus* 6.85-fold change, *P* = 0.72, Table S6). Deletion of *bla*_Mab_ significantly improved the activity of imipenem alone at the two concentrations tested (0.17-fold *versus* 1.75-fold change at 8 μg/mL and 0.14-fold *versus* 0.74-fold change at 32 μg/mL; *P* < 0.05 for both comparisons).

The addition of tedizolid improved the activity of imipenem at 8 and 32 μg/mL against the Δ*bla*_Mab_ derivative leading to 95% and 96% of killing respectively. At a low dose of imipenem, production of Bla_Mab_ moderately reduced the activity of imipenem-tedizolid but the difference was not statistically significant (95% *versus* 90% killing, respectively; *P* = 0.37). This difference was slightly more pronounced for the combination involving the high dose imipenem (96% and 79% killing) reaching statistical significance (*P* < 0.05).

Since production of Bla_Mab_ reduced the efficacy of the imipenem-tedizolid combination, we tested whether inhibition of Bla_Mab_ by avibactam at 16 μg/mL could improve the intramacrophage activity of the combination against *M. abscessus* CIP104536 (Fig. 2B; Table S7). As previously described (16,26), the activity of imipenem at 8 μg/mL was improved by avibactam leading to a 0.32-fold change in the number of CFUs. This fold change is similar to that observed for the imipenem-tedizolid combination (0.28-fold change). The triple combination comprising tedizolid, imipenem, and avibactam was the more active regimens leading to a 97% of killing. Increasing the concentration of imipenem from 8 μg/ml to 32 μg/mL did not improve the activity of the triple combination. In conclusion, the addition of avibactam significantly increased the activity of the imipenem-tedizolid combination if imipenem is used at a low concentration (8 μg/ml).

### Impact of the addition of rifabutin on the intramacrophage activity of tedizolid alone or in combination with imipenem or with imipenem-avibactam

We have recently reported that rifabutin increased the activity of imipenem against *M*. *abscessus* in the macrophage model (26). We have evaluated the benefit of adding rifabutin to the combination imipenem-tedizolid with or without avibactam (Fig. 3, and Table S8). Rifabutin at 1 μg/mL, corresponding to concentration achievable in the serum, significantly improved the activity of the triple combination imipenem-avibactam-tedizolid (0.09-fold *versus* 0.28-fold change; *P* < 0.05). Increasing concentration of rifabutin (8 μg/mL,) did not improve the activity of the quadruple combination (Fig S1).

**FIG 3.**
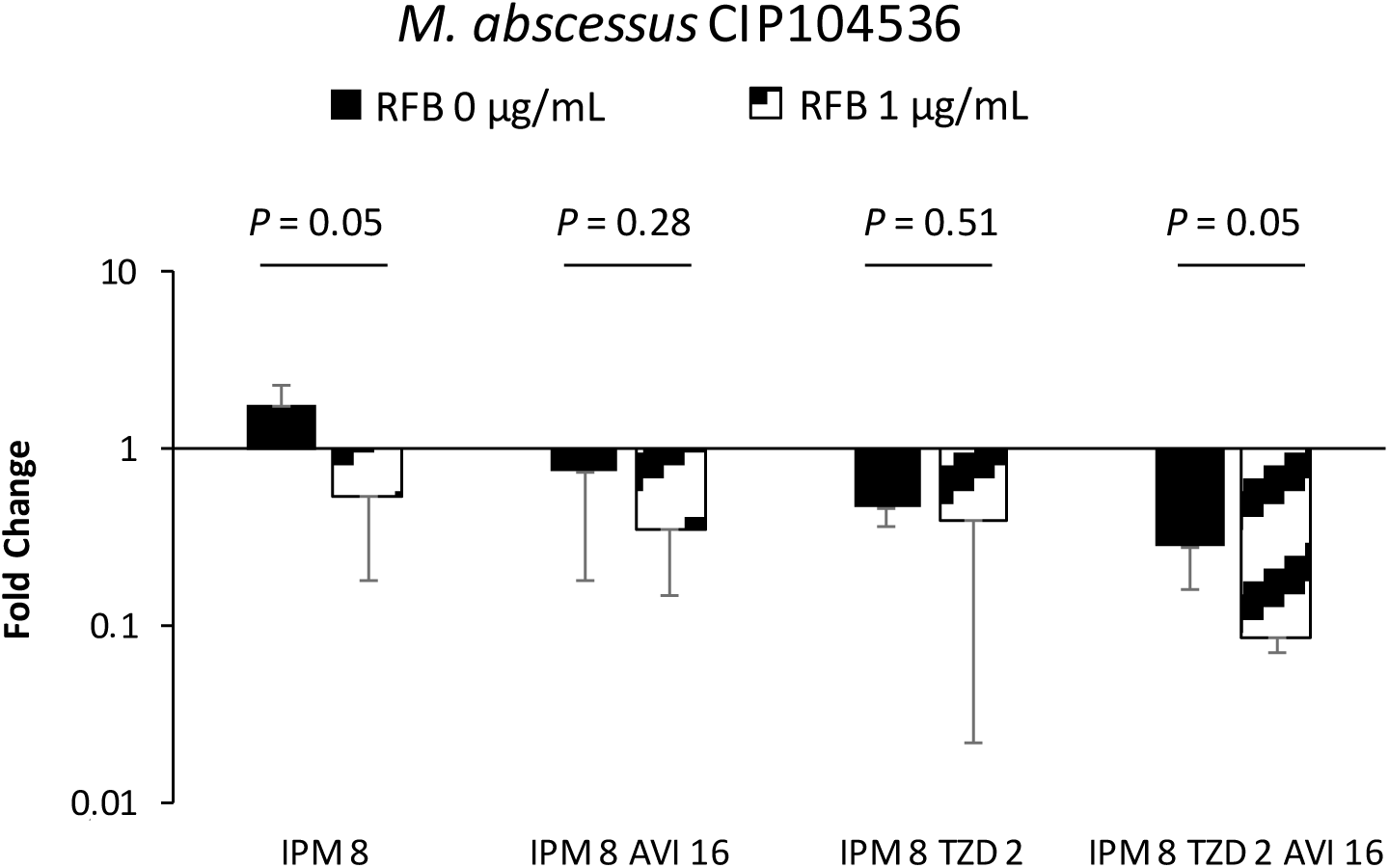
Impact of the addition of rifabutin on the intracellular activity of tedizolid, imipenem, and avibactam against *M. abscessus* CIP104536 in the macrophage model. Rifabutin (RFB) at 1 μg/mL was tested in combination with imipenem (IPM) at 8 μg/mL, tedizolid (TZD) at 2 μg/mL, and avibactam (AVI) at 16 μg/mL.

### Impact of sub-concentration of tedizolid on β-lactamase activity

To explore the mechanism underlying the improved activity of imipenem when combined to tedizolid, we determined the β-lactamase specific activity in crude bacterial extracts using nitrocefin as the substrate (Fig. 4). Growth of *M. abscessus* CIP104536 in presence of sub-concentrations of tedizolid (1 μg/ml and 2 μg/ml, corresponding to ¼ and ½ of the MIC) decreased the β- lactamase specific activity (from 50 to 28 and 14 nmol/min/mg, respectively; *P* < 0.05 for both comparison). These results suggest that a decrease in the production of the β- lactamase Bla_Mab_ could contribute to the antibacterial activity of the tedizolid-imipenem combination.

**FIG 4.**
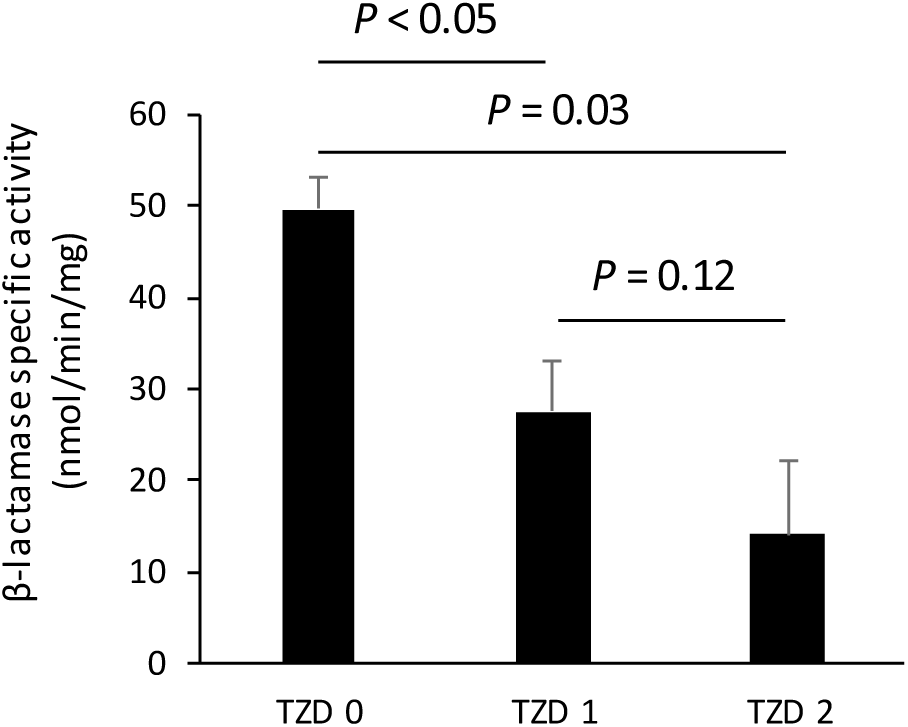
Specific β-lactamase activity in crude extracts from *M. abscessus* CIP104536 grown in 7h9sB in the absence (TZD 0) or presence of sub-inhibitory concentrations of tedizolid (TZD at 1 or 2 μg/mL).

## DISCUSSION

Since tedizolid has recently been reported to display promising *in vitro* activity against *M*. *abscessus* (24), we first investigated the *in vitro* and intra-macrophage activity of combinations comprising this drug and imipenem, which is the recommended β-lactam for the treatment of pulmonary infections in cystic fibrosis patients (9). The activities have been evaluated using imipenem alone or in combination with avibactam, the only approved β- lactamase inhibitor active on Bla_Mab_ (13). The impact of the β-lactamase Bla_Mab_ was also evaluated by comparing *M. abscessus* CIP104536 and its Bla_Mab_-deficient derivative obtained by deletion of the *bla*_Mab_ gene (13,17). Tedizolid was tested at 2 μg/mL corresponding to a therapeutically achievable concentration in serum (27). A 10-to 15-fold intracellular accumulation of tedizolid has been reported in human macrophages (28). The intramacrophage accumulation may account for the observed activity of tedizolid in the macrophage model at a concentration lower than the MIC (4 μg/mL). *In vitro*, the combination of imipenem and tedizolid was moderately synergistic (Table 1) and moderately active in the time kill curve assay (Fig. 2). Inhibition of Bla_Mab_ by avibactam and comparison of *M. abscessus* CIP104536 and its Bla_Mab_-deficient derivative both indicated that hydrolysis of imipenem by Bla_Mab_ had a minor impact on the *in vitro* activity of imipenem. In contrast, Bla_Mab_ significantly impaired the activity of imipenem in the macrophage model. As previously described, this difference may be accounted for by a 20-fold induction of Bla_Mab_ synthesis in the macrophage (16). The combination of tedizolid and imipenem at a low dose (8 μg/mL) was similarly active against *M. abscessus* CIP104536 and its Δ*bla*_Mab_ derivative. This observation suggests that inhibition of the production Bla_Mab_ by tedizolid may contribute to the activity of the tedizolid-imipenem combination in agreement with the negative impact of sub-inhibitory concentrations of tedizolid on β-lactamase production observed *in vitro* (Fig. 4). In the macrophage, the extent of killing of *M. abscessus* CIP104536 by the imipenem-tedizolid combination was further increased by the addition of avibactam. Thus, both tedizolid and avibactam should be considered as potential therapeutic options to improve the activity of imipenem.

Since it has recently been shown that rifabutin has a promising activity against *M. abscessus* (26,29), the second objective of our study was to compare the activity of tedizolid and rifabutin and to evaluate whether combinations comprising these two drugs have a potential therapeutic interest. In the macrophage, tedizolid and rifabutin were similarly active in improving the activity of imipenem alone (Fig. 5). Testing tedizolid and rifabutin (Fig. 3) revealed the benefit of combining these two drugs with imipenem.

**FIG 5.**
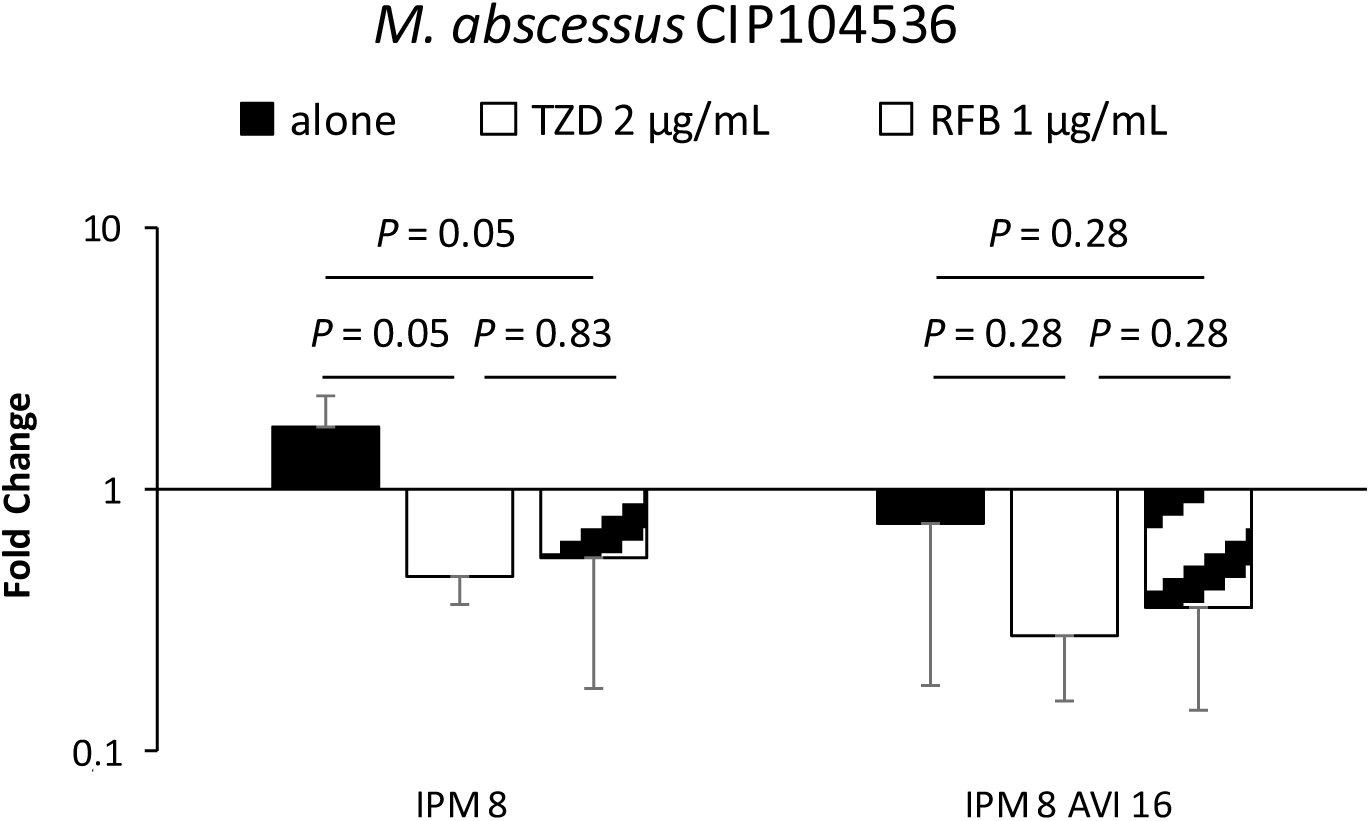
Impact of addition of rifabutin and tedizolid on the intracellular activity of imipenem and avibactam against *M. abscessus* CIP104536 in the macrophage model. Rifabutin (RFB) at 1 μg/mL and tedizolid (TZD) at 2 μg/mL were tested in combination with imipenem (IPM) at 8 μg/mL alone or combined with avibactam (AVI) at 16 μg/mL.

In conclusion, the assessment of the efficacy of drug combinations in the macrophage indicates that imipenem-avibactam-tedizolid-rifabutin should be clinically evaluated, particularly in infections due to macrolide-resistant *M. abscessus*. The advantage of tedizolid and rifabutin as potential therapeutic options for the treatment of lung infections in CF patients is also supported by the relatively low toxicity of these drugs and their oral route of administration. Thus, tedizolid and rifabutin could in particular be considered as alternatives to amikacin in the recommended treatment.

## MATERIAL AND METHODS

### Bacterial strains and growth conditions

*M. abscessus* CIP104536 (ATCC19977) and its β- lactamase-deficient derivative (Δ*bla*_Mab_) (13) were grown in 7H9sB (Middlebrook 7H9 broth (BD-Difco, Le Pont de Claix, France) supplemented with 10% (vol/vol) oleic acid, albumin, dextrose, catalase (OADC; BD-Difco) and 0.05% (vol/vol) Tween 80 (Sigma-Aldrich) at 30°C with shaking (150 rpm) (1).

### Antibiotics

Imipenem, rifabutin, and avibactam were purchased from Mylan (Saint-Priest, France), Serb laboratories (Paris, France), and Advanced ChemBlocks Inc (Burlingame, CA), respectively. Tedizolid was kindly provided by MSD (Kenilworth, USA). Water was the solvent for preparing stock solutions for imipenem and avibactam. Tedizolid was solubilized in dimethylsulfoxyde and rifabutin was solubilized in 96% ethanol.

### MIC determination

MICs were determined in 96-well round-bottom microplates using the microdilution method as previously described (26). The MIC was defined as the lowest drug concentration preventing the resazurin color change from blue to pink or violet. The experiments were performed in quintuplet. Data are the medians from five experiments.

### Synergy testing

Synergistic activities were tested by the two-dimensional dilution checkerboard method as previously described (26) The experiments were performed in quintuplet. According to the American Society for Microbiology, a combination with a FIC index ≤ 0.5 is considered as synergistic.

### Time-kill assays

Time-kill assays were performed as previously described (16, 26). Briefly, Erlen-meyer flasks containing 20 mL of 7H9sB supplemented with imipenem (8 μg/mL), tedizolid (2 μg/mL), and avibactam (4 μg/mL), alone or in combination, were inoculated with exponentially-growing bacteria of *M*. *abscessus* CIP104536 or its Δ*bla*_Mab_ derivative (5 x 10^6^ cfu/mL) and incubated with shaking at 30°C for 72 h. Due to its instability, imipenem (8 μg/mL) was added each 24h. At 0, 48, and 72 h, bacteria were enumerated (colony-forming units, CFUs) and the plates were incubated for 4 days at 30°C. Results are the mean ± standard deviation from three experiments.

### Activity of antibiotics in THP-1 macrophages model

The activity of antibiotics was studied on a THP1 macrophage infection model as previously described (16,26). Briefly, THP-1 cells were seeded into 24-well plates (5 × 10^5^ cells per 1-mL well), differentiated for 24 h, and infected with *M. abscessus* CIP104536 or its Δ*bla*_Mab_ derivative. Imipenem (8 and 32 μg/mL), tedizolid (2 μg/mL), avibactam (16 μg/mL) and rifabutin (1 and 8 μg/mL), alone or in combination, were added to each well. Plates were incubated with 5% CO_2_ at 37°C for 2 days. Imipenem was added each 24 h. The number of CFUs was counted after 4 days of incubation at 30°C by plating serial dilutions of macrophage lysates. Results are the mean ± standard deviation of three experiments.

### Specific β-lactamase activity of total protein extracts

*M. abscessus* was grown in 7h9sB containing sub-inhibitory concentrations of tedizolid (1 and 2 μg/mL), corresponding to ¼ and ½ of the MIC of tedizolid, for 48 hours at 37°C. Bacteria were collected by centrifugation, resuspended in 2 mM piperazine-N,N’-bis(2-ethanesulfonic acid (pH 6.5; PIPES), and lyzed with glass beads (Fast Prep; Thermosavant). Proteins were determined by the Bradford method. β-lactamase activity was determined at 20°C by spectrophotometry at 480 nm in 2 mM PIPES (pH 6.5) using nitrocefin (100 μM) as the substrate (Carry-100 Bio spectrophotometer; Varian). The results are the mean ± standard deviation from three experiments.

### Statistical analysis

The comparison of the antibiotic activities was done using The Mann-Whitney U test. Statistical analyses were performed with EPI Info(tm) software version 7.1.3 (Center for Disease Control and prevention, Atlanta).

## Transparency declaration

J.-L. M. has received consulting fees (scientific advisor for ceftolozane-tazobactam, Merck Sharp and Dohme) and reimbursement of travel expenses (attendance at 54^th^ Interscience Conference on Antimicrobial Agents and Chemotherapy, 2014 and attendance at 27^th^ European Congress of Clinical Microbiology and Infectious Diseases, 2017) from AstraZeneca and Merck Sharp and Dohme, respectively. All other authors: none to declare.

